# Evaluation of neutralizing antibodies produced by papaya mosaic virus nanoparticles fused to the E2EP3 peptide epitope of Chikungunya envelope

**DOI:** 10.1101/2020.02.28.970657

**Authors:** Hussin A. Rothan, Rohana Yusof

## Abstract

Chikungunya virus (CHIKV) infection is the cause of acute symptoms and chronic symmetrical polyarthritis associated with long-term morbidity and mortality. Currently, there is no available licensed vaccine or particularly useful drug for human use against CHIKV infection. This study was conducted to evaluate the efficacy of antibodies produced by papaya mosaic virus (PapMV) nanoparticles fused to E2EP3 peptide of CHIKV envelope as a recombinant CHIKV vaccine. PapMV, PapMV E2EP3, and E2EP3 PapMV were produced in E. coli with an approximate size of 27 to 30 kDa. ICR mice (5 to 6 weeks of age) were injected subcutaneously with 25 micrograms of vaccine construct, and ELISA measured the titer of CHIKV specific IgG antibodies. The results showed that both recombinant proteins E2EP3 PapMV and PapMV E2EP3 were able to induce IgG antibodies production in immunized mice against CHIKV while immunization with recombinant PapMV showed no IgG antibodies induction. The neutralizing activity of the antibodies generated by either E2EP3 PapMV or PapMV E2EP3 exhibited similar inhibition to CHIKV replication in Vero cells using the cells based antibody neutralizing assay and analyzed by plaque formation assay. This study showed the effectiveness of nanoparticles vaccine generated by fusing epitope peptide of CHIKV envelope to papaya mosaic virus envelope in inducing a robust immune response in mice against CHIKV. The data showed that levels of neutralizing antibodies correlate with a protective immune response CHIKV replication.

## Introduction

Chikungunya virus (CHIKV) is a mosquito-borne enveloped alphavirus belong to the family of *Togaviridae.* CHIKV contains a single-stranded positive-sense RNA genome of approximately 11.8 kbp, which is capped and polyadenylated, encoding two open reading frames (ORFs)[1]. The 5 ORF is expressed via cap-dependent translation as an ns-P1-3 or ns-P1-4 polyprotein, and it is cleaved by an nsP2 encoded protease. The structural protein ORF is translated into three main structural proteins: the capsid (C), the envelope glycoproteins E2 and E1, and two other proteins of E3 and 6K[2]. The envelop polyprotein of E3E26KE1 is then processed in the endoplasmic reticulum. The mature virion contains 240 of E2/E1 heterodimers arranged in 80 trimeric spikes on the surface, which are inserted into the plasma membrane of infected cells after transport through the secretory pathway. The E2 trimeric spikes are essential for budding of new virus particles and host receptor recognition and attachment, while E1 is responsible for cell entry via pH-dependent endocytosis. The capsid protein is cleaved from the structural polyprotein and encapsidates the cytoplasmic viral genomic RNA. Cytoplasmic nucleocapsids containing the genomic RNA and 240 copies of the capsid protein bud from the cells surface to acquire the virion envelope and envelope protein spikes[3,4].

At present, there is no available licensed vaccine or particularly useful drug for human use for any *alphavirus.* Several pre-clinical CHIKV vaccines were described. Inactivated virus formulation, like a formalin-inactivated *alphavirus* vaccine, has proved to be immunogenic in humans [5,6]. Live-attenuated CHIKV vaccines are found to be immunogenic, but they have side effects like arthralgia in a human phase II study [7,8]. Antibodies are required for protection against CHIKV infections, but DNA vaccines have not been able to generate antibody responses in humans[6,9] effectively. Chimeric virus vaccines, recombinant adenovirus vaccines, subunit protein vaccines are other examples of CHIKV vaccines[10,11]. In this study, an 18-amino acid peptide derived from CHIKV envelop was fused to the Papaya Mosaic virus envelop (PapMV) as a platform for creating vaccine nanoparticles. The recombinant vaccine was able to induce an immune response in mice against CHIKV. The data showed that levels of neutralizing antibodies correlate with a protective immune response, which can accelerate the development accessibility of CHIKV.

## Methods

### 1. Virus and cells

A clinical isolate of CHIKV recovered from the serum of infected patients (CHIKV isolate, SGEHICHS277108, Accession FJ445510) was used for propagation by a single passage in C6/36 mosquito cells. Vero cells (ATCC CCL-81) were obtained from the American Type Culture Collection (ATCC; Rockville, MD, USA) and cultured in complete Dulbecco’s modified Eagle’s medium (DMEM) supplemented with 10% fetal bovine serum (FBS) as growth medium or 2% FBS as maintenance medium.

### 2. Cloning and expression of CHIKV nanoparticles vaccine

Recombinant Papaya virus particles fused with CHIKV peptide epitope E2EP3. Recombinant plasmid construction was carried out as previously described[12–14]. In brief, protein sequences of PapMV, PapMV-E2EP3, and E2EP3-PapMV were converted to DNA sequences and optimized based on *E. coli* preferred codons. Oligonucleotides of 60 mer were designed with an overlapping 3’ ends (15 nucleotides) using the Primer3 (version 0.4.0) analysis software. The long oligonucleotides were annealed, and the mixture of the complementary overlapping oligonucleotides was extended and filled-in with dNTP and Klenow DNA polymerase (Invitrogen, USA). The full DNA fragment was cloned downstream of T7 promoter of *the E. coli* expression vector. The recombinant plasmid DNA was transformed into E. coli BL21, followed by screening and restriction digestion. The recombinant plasmids were isolated, and the purified plasmids were sequenced to verify the correct sequences without mutations. *E. coli* harboring recombinant vectors were cultured in 1 L of LB broth medium supplemented with ampicillin and incubated at 37°C. The bacterial cells were harvested, lysed, and sonicated on the ice after the induction of protein expression by IPTG. Recombinant protein in the supernatant was purified to 90% purity by a single-step chromatography on Ni^++^ - metal chelate affinity (Qiagen, UK) columns and eluting at 0.5 M imidazole. The eluted protein profile was analyzed using SDS-PAGE, and the peak fractions were pooled[12,15]. The purified proteins were concentrated via Vivaspin^®^ 20 concentrator tubes (Viva products, Littleton, MA, USA).

### 3. Immunization

*In vivo* experiments were carried out following the University of Malaya guidelines on the Care and Use of Laboratory Animals following approval of the animal ethics protocols used in the investigation by the Animal Ethics Committee of the University of Malaya. Twelve ICR mice (5–6 weeks of age) were injected subcutaneously with 25 μg of PapMV, E2EP3-PapMV, and PapMV-E2EP3 or endotoxin-free PBS (each group 3 mice). Primary immunization was followed by one booster dose given two weeks later. Sera samples were collected after 21 days after the first injection and stored at −20 °C until analysis.

### 4. Determination of the CHIKV-specific IgG antibodies

The titer of CHIKV-specific IgG antibodies was performed by ELISA, as previously described[16]. In brief, 96-well plates were coated with purified CHIKV antigen in PBS for overnight, blocked with blocking buffer (PBS containing 0.05% Tween 20 and 5% ABS) and incubated for two h at room temperature. Plasma samples were then diluted at 1:100 to 1:10,000 in blocking buffer and incubated for two h at room temperature. HRP-conjugated goat anti-mouse IgG was used to detect mouse antibodies bound to virus-coated wells. Reaction mixtures were developed using TMB (3,3,5,5-tetramethylbenzidine) substrate (Sigma-Aldrich), and the absorbance was measured at 450 nm.

### 5. In vitro neutralization

Neutralizing activity of antibodies was tested in triplicate and analyzed by plaque formation assay in Vero cells. CHIKV was mixed at a multiplicity of infection (MOI) of 5, with either heat-inactivated mice serum (between 1:50and 1:10,000) and incubated for 2 h at 37°C, with gentle agitation. Virusantibody mixtures were then added to Vero cells seeded into 6-well plates and incubated for 2 h at 37°C with gentle shaking every 15 min. The medium was removed, and cells were washed with PBS and a new DMEM medium supplied with 2% FBS and incubated for 72 h at 37°C. Culture supernatant was collected, and virus titers were determined by plaque formation assay.

### 6. Plaque formation assay

This assay was carried out as previously described[17–19]. In brief, A 10-fold serial dilution of culture supernatant of CHIKV infected cells was added to fresh Vero cells grown in 6-well plates (0.5 × 10^6^ cells) and incubated for 1 h at 37°C. The cells were overlaid with DMEM (maintenance medium) containing 0.5% agarose. Viral plaques were stained with crystal violet dye after 5-day of incubation.

Virus titers were calculated according to the following formula: Titer (p.f.u./ml) = number of plaques/volume of the diluted virus added to the well × dilution factor of the virus used to infect the well in which the plaques were enumerated.

### 7. Statistical analysis

All the assays were performed in quadruplicate, and the statistical analyses were performed using GraphPad Prism version 5.01 (GraphPad Software, San Diego, CA). P values <0.05 were considered significant. The error bars are expressed as ± SD.

## Results and Discussion

CHIKV disease is characterized by acute symptoms that generally last about a week and are selflimiting and chronic symmetrical polyarthritis/polyarthralgia lasting for months to years, indicating long-term morbidity and mortality of CHIKV[20–22]. In recent years, CHIKV disease has become a global public health problem. However, there is no licensed vaccine available for CHIKV. The current study provides a method of producing recombinant CHIKV vaccine nanoparticles.

### 1. Production of the recombinant vaccine nanoparticles

In this study, an 18-amino acid peptide (E2EP3_2800-2818_) derived from CHIKV envelope was fused to the Papaya Mosaic virus envelop (PapMV) as a platform for creating vaccine nanoparticles. Five residues extension from N- and C-termini of E2EP3 peptide were included for favor natural processing of the peptide. The position of E2EP3 peptide might influence the formation and immunogenicity of the nanoparticles. Thus, the peptide was fused to either N- or C-termini of PapMV (Fig. 1A). To achieve this aim, three recombinant plasmid constructs were generated to express the Papaya Mosaic virus envelop without insertion *(PapMV)*, with insertion at the N-terminus *(E2EP3-PapMV)* and with insertion at C-terminus *(PapMV-E2EP3)* as presented in Figure 1A. The recombinant plasmids were transformed into *E. coli* to express three recombinant proteins named PapMV, E2EP3-PapMV, and PapMV-E2EP3[23]. The recombinant proteins were affinity purified on a Ni^++^-column via 6XHis tag. The expected size of these proteins was approximately 27 to 30 kD. (Fig. 1B). The concentrated recombinant proteins were visualized under transmission electron microscopy. The results obtained by transmission electron microscopy showed there are two structures of nanoparticles, typical long rod-shaped (approximately 120 nm in length) and discshaped structure (approximately 25 nm diameter) for PapMV, E2EP3-PapMV and PapMV-E2EP3 (Fig. 1C).

**Figure 1.**
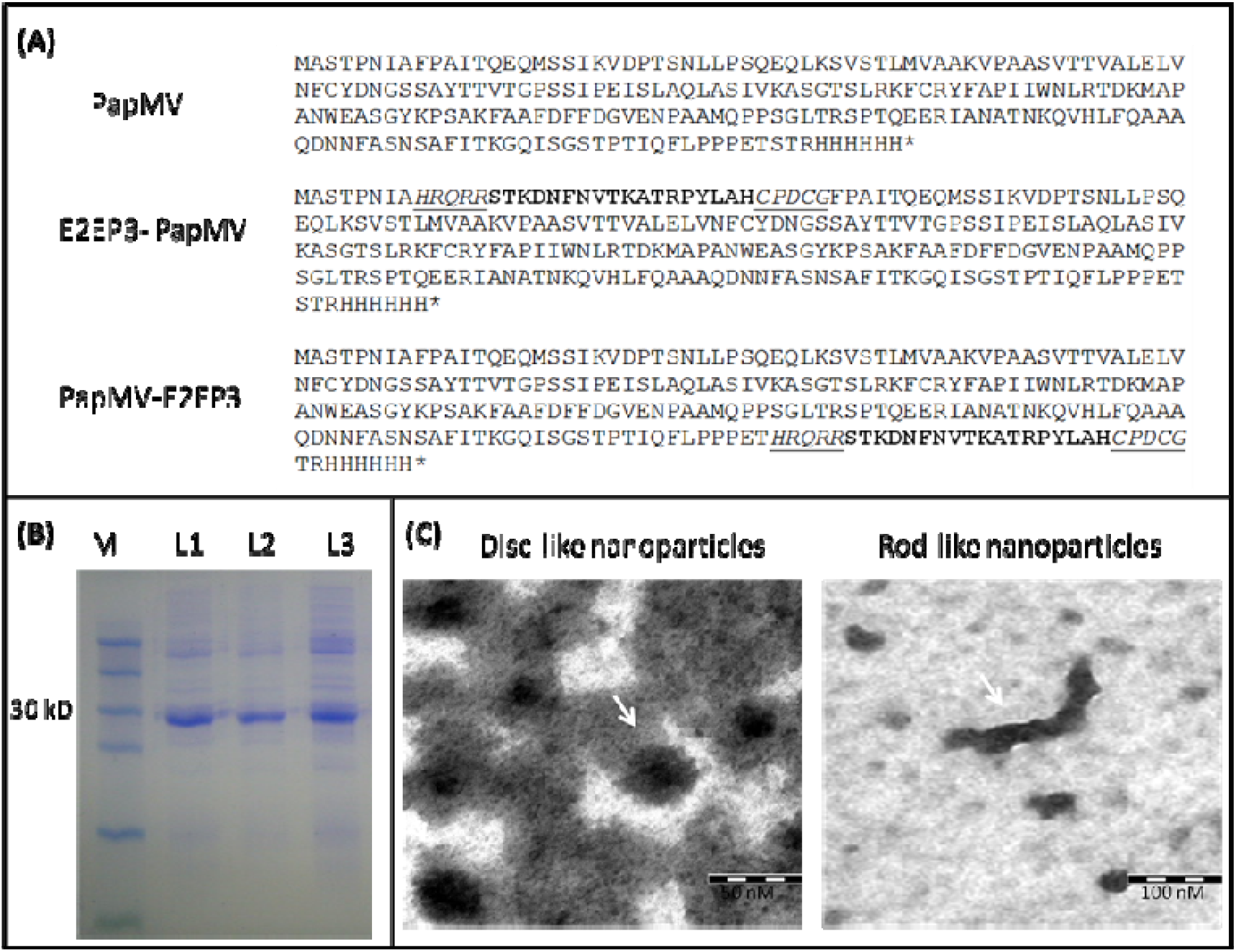
Design and expression of recombinant CHIKV vaccine nanoparticles. **(A)** The amino acid sequence of E2EP3 peptide derived from CHIKV envelop was fused to the Papaya Mosaic virus envelop (PapMV) as a platform for creating vaccine nanoparticles. The peptide sequence (bold) was flanked from N- and C-termini with five residues extension from CHIKV envelop (italic-underlined) for favor natural processing of the peptide. The peptide was fused either at N- or C-terminus resulting in two recombinant proteins PapMV, E2EP3-PapMV and PapMV-E2EP3 or without insertion PapMV. **(B)** The recombinant proteins were affinity purified on a Ni^++^-column via 6XHis tag. The expected size of these proteins was approximately 27 to 30 kD based on SDS-page results, M (marker), L1 (PapMV), L2 (E2EP3-PapMV) and L3 (PapMV-E2EP3). **(C)** The transmission electron microscopy showed there are two structures of nanoparticles, a disc-shaped structure (approximately 25 nm diameter) and long rod-shaped (about 120 nm in length) for PapMV, E2EP3-PapMV and PapMV-E2EP3.

### 2. The recombinant nanoparticles induced an immune response in mice

In this study, the immune response to PapMV nanoparticles was examined by a subcutaneous injection of twelve mice with 25 μg each of the recombinant PapMV, E2EP3-PapMV, PapMV-E2EP3 or endotoxin-free PBS. A booster dose was given on day 15 after primary immunization. Mice sera were assayed for anti-PapMV, E2EP3-PapMV, and PapMV-E2EP3 antibodies. The results showed both recombinant proteins E2EP3-PapMV and PapMV-E2EP3 were able to induce IgG antibodies production in immunized mice against CHIKV while immunization with recombinant PapMV or endotoxin-free PBS showed no IgG antibodies induction (Fig. 2). Our data are supported by recent studies that show that protective antibodies play a role in the control of CHIKV infection. Previous studies show that the neutralization of viruses by anti-CHIKV antibodies protects against CHIK infection[4,24]. Natural CHIKV infections induce high neutralizing antibody levels from 40 to 20,000 in titter[25], which have been detected in CHIKV patients during the infection course. The presence of IgG antibodies correlates with viral clearance and long-term protection[26].

**Figure 2.**
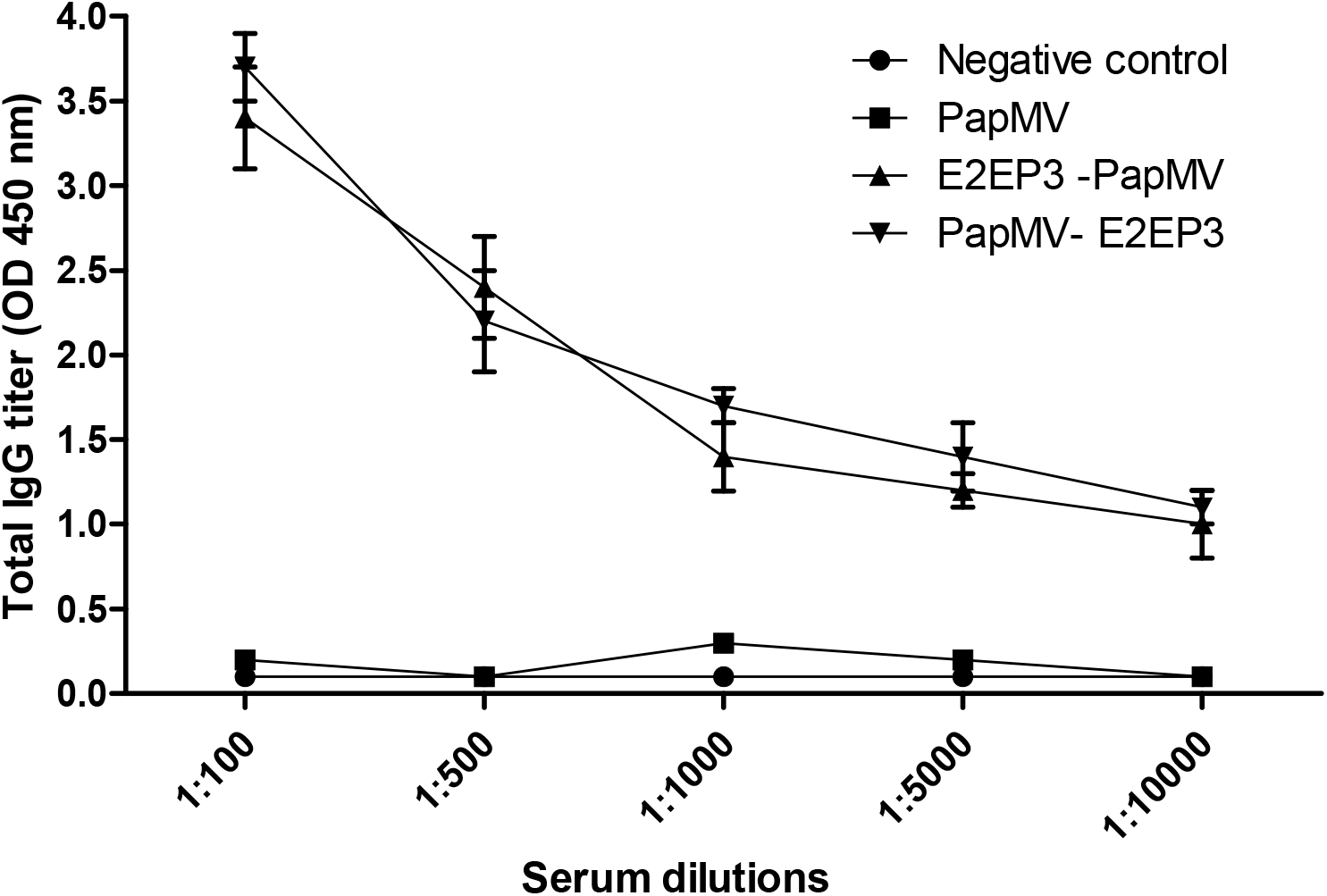
Antibody profiles of PapMV nanoparticles in immunized mice. Virus-specific IgG antibody titers in each serum sample were determined by ELISA using purified CHIKV virions. Mice were injected subcutaneously with 25 μg of PapMV, E2EP3-PapMV, and PapMV-E2EP3 or endotoxin-free PBS. Primary immunization was followed by one booster dose given 2 weeks later. Sera samples were collected 21 days post-injection and diluted at 1:100 to 1:10,000 in blocking buffer for ELISA assay. The recombinant proteins E2EP3-PapMV and PapMV-E2EP3 were able to induce IgG antibodies production in immunized mice against CHIKV while immunization with recombinant PapMV or endotoxin-free PBS showed no IgG antibodies induction.

### 3. The sera of immunized mice inhibited virus infection in vitro

One of the advantages of peptidic vaccination is the good mimicry of the fused epitope. To check if the peptide derived from the CHIKV surface glycoprotein that is fused to the C-terminal or N-terminal of the PapMV can exhibit differences antibodies affinity to CHIKV. The neutralizing activity of antibodies generated by either E2EP3-PapMV or PapMV-E2EP3 was tested in vitro using the cells-based antibody neutralizing assay and analyzed by plaque formation assay. CHIKV was mixed at a multiplicity of infection (MOI) of 5, with either heat-inactivated mice serum using increasing dilution factor. The results showed that E2EP3-PapMV and PapMV-E2EP3 exhibited similar inhibition to CHIKV replication in Vero cells (Fig. 3A and 3B). Dilutions at 1:50 to 1:200 of mice sera containing anti-CHIKV antibodies were able to neutralize more than 80% of CHIKV replication compare to the higher dilution factors.

**Figure 3.**
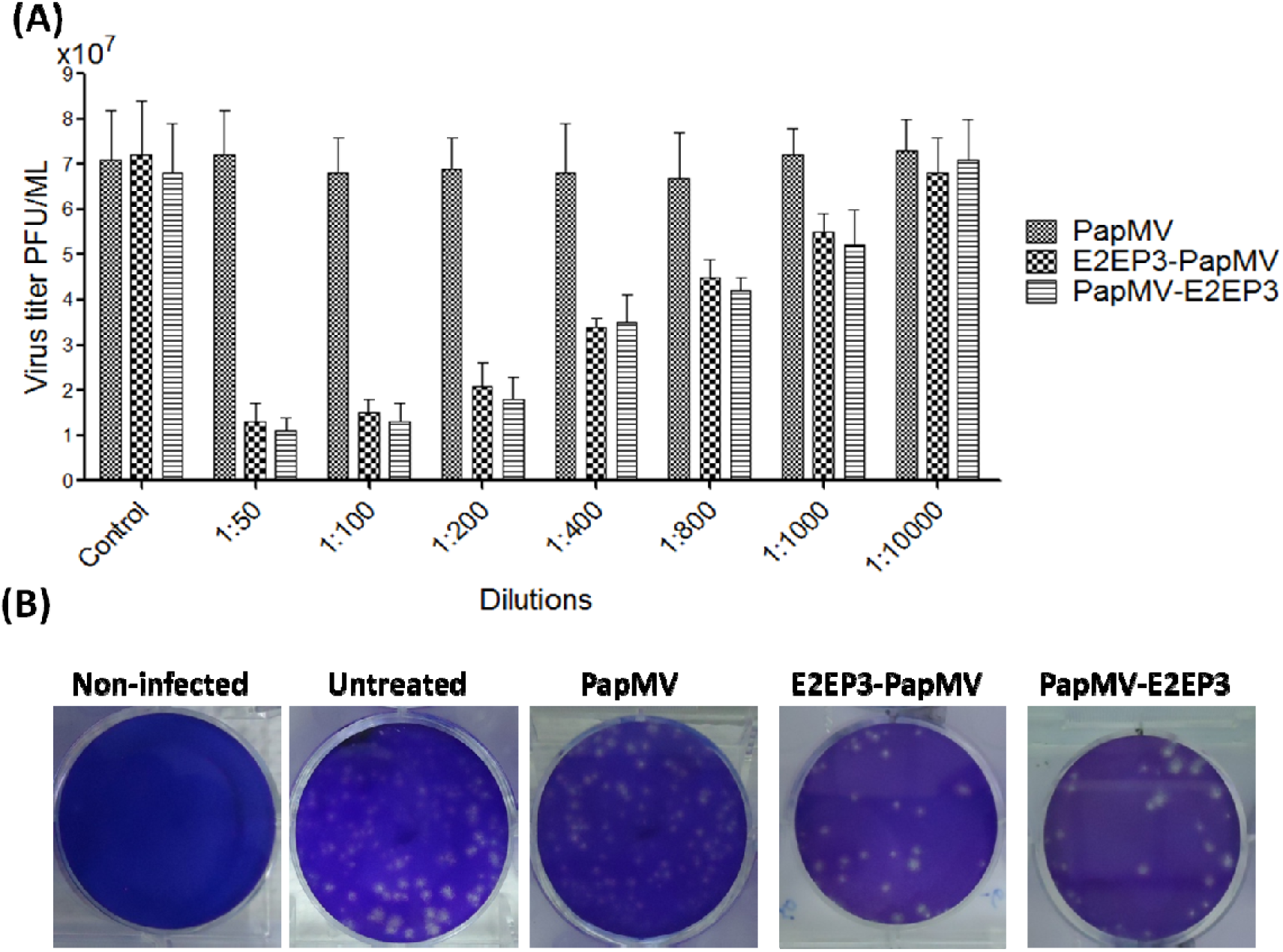
Evaluation of neutralizing activity of mice sera containing antibodies against CHIKV replication in Vero cells. **(A)** CHIKV was mixed with heat-inactivated mice sera at serial dilutions ranged from 1:50 to 1:10,000. After incubation for 2 h at 37°C, virus-antibody mixtures were then added to Vero cells at an MOI of 5 and incubated for 72 h. Culture supernatant was collected and virus titers were determined by plaque formation assay. **(B)** A serial dilution of culture supernatant of CHIKV infected cells treated with 1:50 dilution was added to fresh Vero cells grown in 6-well plates. The cells were overlaid with DMEM and viral plaques were stained with crystal violet dye after 5-day of incubation.

It has been known that Virus-like particle (VLP) formulation mimics the native virus surface architecture and protein conformation, and it induces protective antibody responses in the absence of adjuvant [27]. A recent CHIKV VLP-based vaccine provides protection in both mice and non-human primates by DNA transfection of mammalian cells, but this system out-performs cost and scalability[28–30], therefore, the CHIKV VLPs in the presence of adjuvant are economical and easy to scale up for an epidemic response. It has been shown that the E2 domain of the CHIKV envelope glycoprotein, which binds with high affinity to CHIKV, is responsible for virus neutralization. This data shows that levels of neutralizing antibodies correlate with a protective immune response, which can accelerate the development accessibility of the CHIKV vaccine[4]. Based on the advantage of nanoparticle vaccine, we sought to test the protection efficiency, and immunogenicity of engineered papaya mosaic virus nanoparticles (PapMV-VLPs) fused with peptide-epitope derived from Chikungunya E2 glycoprotein to trigger a protective immune response against CHIKV. The PapMV platform is rod-shaped (15nm in diameter~150-250nm in length, which was recently shown to induce a high antibody response against a hepatitis C virus (HCV) epitope[31,32].

In conclusion, vaccination is the most cost-effective means of protecting the at-risk populations in CHIK-endemic developing countries. CHIK epidemics are, however, explosive and rapidly moving, but not predictable. This study showed the effectiveness of nanoparticles vaccine generated by fusing epitope peptide of CHIKV envelope to papaya mosaic virus envelop (PapMV). The recombinant vaccine was able to induce an immune response in mice against CHIKV. The data showed that levels of neutralizing antibodies correlate with a protective immune response, which can accelerate the development accessibility of CHIKV.

#### Ethical approval

Animal experiment was carried out following the University of Malaya guidelines on the Care and Use of Laboratory Animals following approval of the animal ethics protocols used in the investigation by the Animal Ethics Committee of the University of Malaya.

